# Zebrafish live imaging reveals a surprisingly small percentage of spinal cord motor neurons die during early development

**DOI:** 10.1101/2024.01.26.577434

**Authors:** Hao Jia, Hongmei Yang, Kathy Qian Luo

## Abstract

It is widely accepted that large numbers of neurons die during the early development of vertebrates; however, the tracking of this dying process in live animals remains challenging. Here, we generated sensor zebrafish achieving live imaging of motor neuron apoptosis at single-cell resolution. Using these sensor zebrafish, we observed for the first time that in an apoptotic motor neuron, caspase-3 activation occurred quickly within 5-6 min and at the same time between the cell body and axon. Interestingly, we found that only a surprisingly small percentage of spinal cord motor neurons died during zebrafish early development, which is quite different from the generally believed massive motor neuron death occurred in the embryonic stage of chicks, mice, rats, and humans. We also observed that most of the apoptotic bodies of dead motor neurons were not colocalized with macrophages. These sensor zebrafish can serve as powerful tools to study motor neuron apoptosis *in vivo*.

## Introduction

Programmed cell death or apoptosis is a dynamic process that occurs in most tissues in live animals. In the nervous system, apoptosis plays an essential role in the regulation of neuronal cell numbers (Hollville, Romero, & Deshmukh, 2019). During the embryonic development of vertebrates, including chicks, mice, rats, and humans, a large number of neuronal cells die through apoptosis (Fuchs & Steller, 2011). In pathological states, such as Alzheimer’s disease, Parkinson’s disease, and motor neuron disease, the death of neuronal cells severely impairs the normal function of the nervous system (Butterfield & Halliwell, 2019; Michel, Hirsch, & Hunot, 2016; Moujalled, Strasser, & Liddell, 2021).

When apoptosis occurs, either through intrinsic or extrinsic pathways, it ultimately leads to the cascade activation of caspases and the degradation of cellular structures, which is no exception for neuronal cells (Blanquie et al., 2017; Heck et al., 2008; Hernandez-Baltazar, Mendoza-Garrido, & Martinez-Fong, 2013). Due to the unique and complex morphology of neuronal cells, especially long axons, many important issues related to neuronal cell apoptosis have not been fully addressed, including: (1) How to visualize neuronal cell death in live animals? (2) How fast caspase-3 can be activated within a single neuron during apoptosis? (3) Which part of a neuron (cell body vs. axon) will degrade first? To address these important questions, we need to develop novel tools that can track neuronal cell apoptosis in live animals.

Previously, we have developed a fluorescence resonance energy transfer (FRET)-based biosensor, named sensor C3, for detecting cancer cell apoptosis. Our *in vitro* results showed that sensor C3 could detect the apoptosis of cancer cells both in 2D and 3D cultures by changing its color from green to blue (Anand, Fu, Teoh, & Luo, 2015; Fu, Peh, Ngan, Wei, & Luo, 2018; Hao, Huang, Wu, Peng, & Luo, 2023; K. Li, Wu, Zhou, Tong, & Luo, 2021; Luo, Vivian, Pu, & Chang, 2001; Tian, Ip, Luo, Chang, & Luo, 2007; Wu, Li, Yuan, & Luo, 2021; Yang, Jia, Zhao, & Luo, 2022). Recently, we showed that sensor C3 could also be used to indicate zebrafish skin cell apoptosis (Jia, Song, Huang, Ge, & Luo, 2020).

In the present study, using *Tol2* transposon-based transgenic technology, we generated novel sensor zebrafish and achieved the noninvasive tracking of motor neuron apoptosis at single-cell resolution in live zebrafish. More importantly, using these sensor zebrafish, we were able to obtain novel insights into the spatiotemporal properties and occurring rates of motor neuron death during early development.

## Results

### Generation of transgenic zebrafish expressing sensor C3 in motor neurons

Sensor C3 is a fusion protein of cyan fluorescent protein (CFP) and yellow fluorescent protein (YFP), between which is an amino acid linker containing the caspase-3 cleavage site DEVD (Figure 1A). In live cells, when sensor C3 is excited with a 458 nm laser, due to the energy transfer between CFP and YFP, the emission light of CFP can excite the YFP to emit green fluorescence, which can be detected at the wavelength of 535 nm. In apoptotic cells, activated caspase-3 proteins cleave sensor C3 at the DEVD site; due to the disruption of the FRET effect, sensor C3 emits blue fluorescence, which can be detected at the wavelength of 480 nm (Figure 1B).

**Figure 1.**
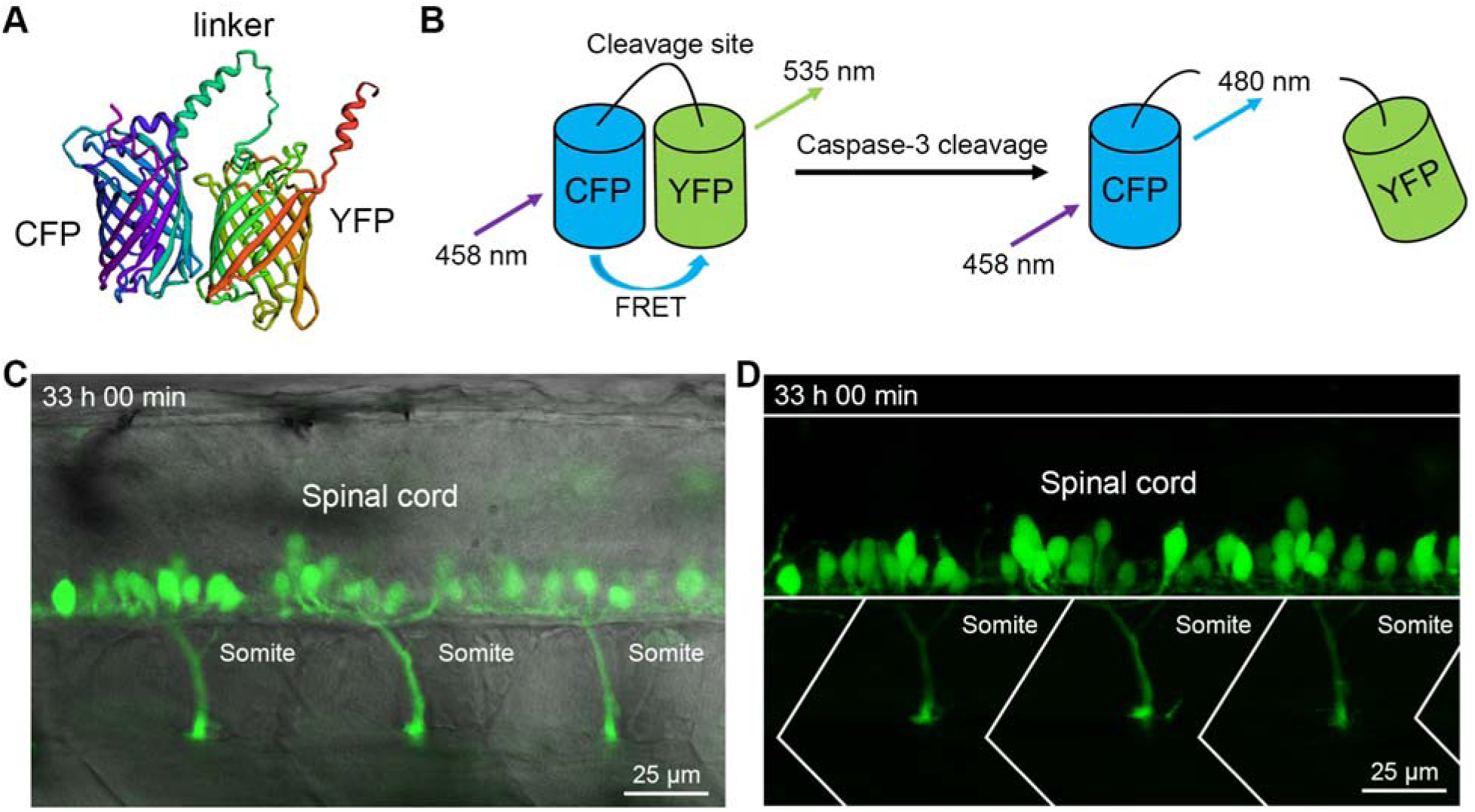
The live imaging of motor neurons in sensor zebrafish. (**A**) The structure of sensor C3 was predicted by using Rosetta software. (**B**) The principle of sensor C3 for apoptosis detection. (**C** and **D**) The expression of sensor C3 in the motor neurons of Tg(*mnx1*:sensor C3) zebrafish. The bright field image and fluorescent image were merged in (C) to show the location of motor neurons and their axons. The anatomical structures and developmental time are indicated.

In our previous publications, we have conducted many experiments to validate the specificity and sensitivity of sensor C3 for detecting caspase-3 activation and apoptosis. First, we confirmed that there was a strong energy transfer from CFP to YFP in purified sensor C3 proteins when CFP was excited. Second, we showed that the cleavage to sensor C3 is specific to caspase-3 and requires the presence of the cleavage sequence of DEVD in sensor C3 proteins. Third, sensor C3 is highly sensitive to caspase-3 cleavage, as it could detect caspase-3 activation at a nanomolar concentration and in 5 min when sensor C3-labelled HeLa cells underwent UV irradiation-induced apoptosis (Luo et al., 2001).

We have confirmed that the apoptosis detected by sensor C3 was consistent with classical apoptotic assays, including chromatin condensation staining, caspase-3 activity assay, DNA fragmentation assay, and morphological changes such as cell shrinkage (Luo et al., 2001; Tian et al., 2007). Furthermore, we showed that sensor C3 could detect the apoptosis of various cancer cells and zebrafish cells by color change no matter whether the apoptosis was induced by UV irradiation, chemical drugs, fluidic shear stress, immune cells-mediated killing, or during normal development (Hao et al., 2023; Jia & Luo, 2021; Jia et al., 2020; K. Li et al., 2021; Luo et al., 2001; Tian et al., 2007; Yang et al., 2022). All aforementioned results show that sensor C3 is a very good tool for detecting apoptosis in live cells and animals.

To detect the death of motor neurons in live animals, we cloned sensor C3 under a motor neuron-specific promoter (Arkhipova et al., 2012; Flanagan-Steet, Fox, Meyer, & Sanes, 2005; Jao, Appel, & Wente, 2012; Rosenberg, Wolman, Franzini-Armstrong, & Granato, 2012; Stil & Drapeau, 2016) and used this construct to generate a transgenic line of sensor zebrafish Tg(*mnx1*:sensor C3) in which the motor neurons specifically expressed sensor C3 proteins. After obtaining the sensor zebrafish, we first examined the morphology of motor neurons in zebrafish. At 33 hours post fertilization (hpf), we observed many green motor neurons inside the spinal cord (Figure 1C, D).

### Visualizing caspase-3 activation and single motor neuron apoptosis during zebrafish early development

To determine the apoptotic status of motor neurons during the development, we took the images of CFP and YFP separately from the Tg(*mnx1*:sensor C3) zebrafish and merged them into FRET images. The green color will indicate the motor neurons are alive, and the blue color will indicate the neurons are apoptotic. The FRET imaging analysis showed that around 30 hpf during zebrafish early development, the cell bodies of some motor neurons appeared in blue. In contrast, most of the cell bodies of these motor neurons appeared in green. Furthermore, the blue signals were also observed in the axon bundle of the motor neurons (Figure 2A-D). Figure 2C showed a motor neuron cluster inside of which more than one motor neuron died. Image stacks of the apoptotic motor neuron in Figure 2D at different optical section levels better illustrated the blue signals in the axon region (Figure 2E). This observation indicates that the majority of the motor neurons are alive, and the minority can undergo apoptosis.

**Figure 2.**
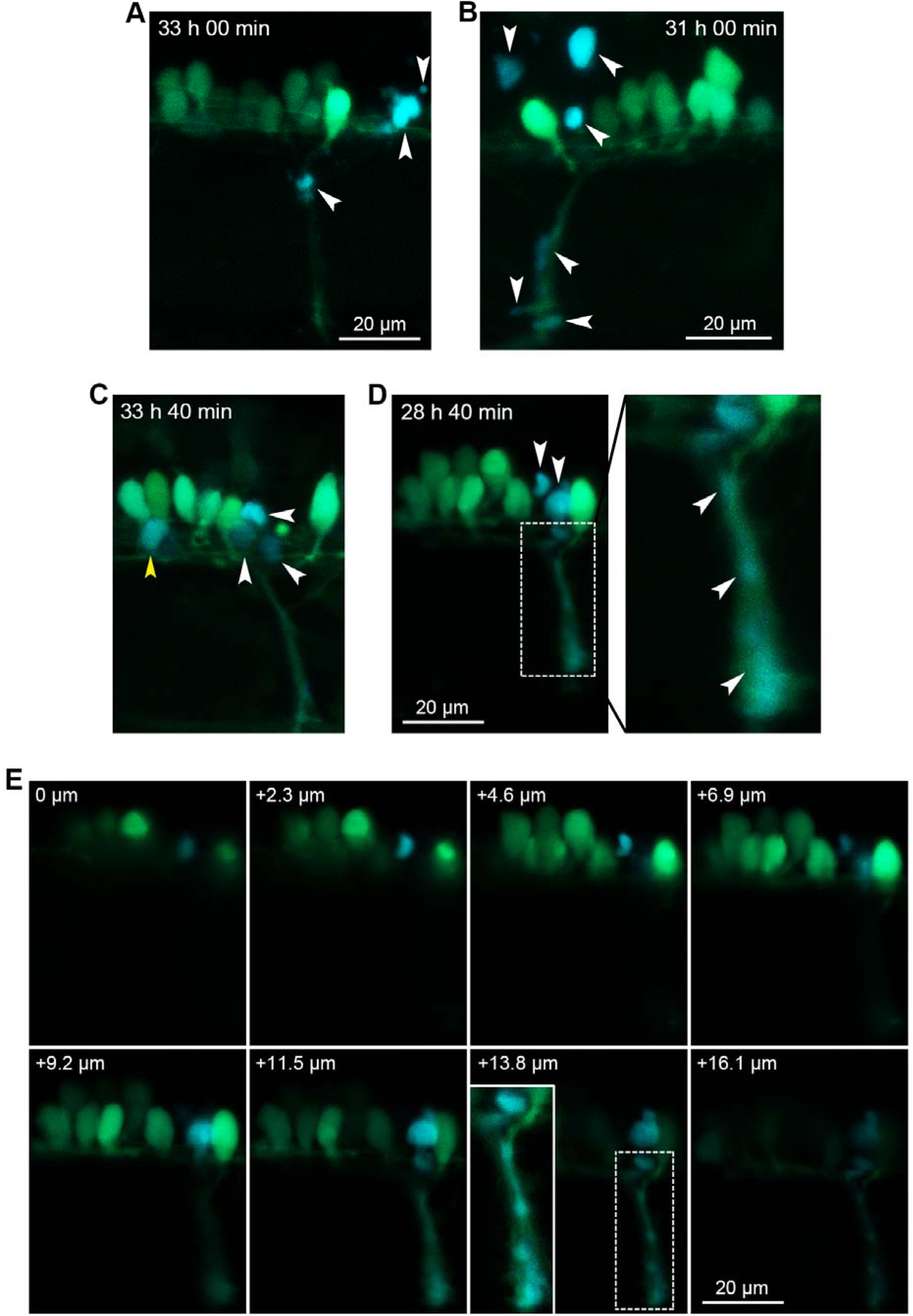
Visualizing caspase-3 activation and single motor neuron apoptosis during zebrafish early development. (**A**−**D**) FRET images of four somites showing motor neurons underwent apoptosis during the development of Tg(*mnx1*:sensor C3) zebrafish. Two dead motor neurons in (C) are indicated with yellow and white arrowheads, respectively. The axon in (D) is enlarged for a better illustration. Blue apoptotic signals from cell bodies and axons are indicated with arrowheads. The developmental time is indicated in each image. (**E**) Z-stack imaging showed the apoptosis of the motor neuron in (D). The axon region in the stack of 13.8 µm is enlarged for a better illustration. The depth of each stack is indicated in each image.

### The caspase-3 activation occurred quickly and almost at the same time between the cell body and axon in apoptotic motor neurons

Images in Figure 2 showed that blue signals appeared in both the cell body and axon of apoptotic motor neurons. To determine which part of a motor neuron degraded first and how fast the degradation process was, we performed time-lapse imaging of caspase-3 activation in live zebrafish. FRET images in Figure 3A showed that at 32 h 15 min of zebrafish development, both the cell body (indicated with a white arrowhead) and the axon (indicated with a yellow arrowhead) of the indicated motor neuron were green, suggesting that caspase-3 was not activated. Just 3 min later, both the cell body and the axon started to change color (32 h 18 min). Another 3 min later, both the cell body and the axon became blue (32 h 21 min). At 32 h 27 min, two blue apoptotic bodies could be observed in the cell body region, and multiple apoptotic bodies were also observed in the axon region. In the next 12 min (until 32 h 39 min), this apoptotic motor neuron gradually degraded into more and smaller apoptotic bodies (Figure 3A; Video S1).

**Figure 3.**
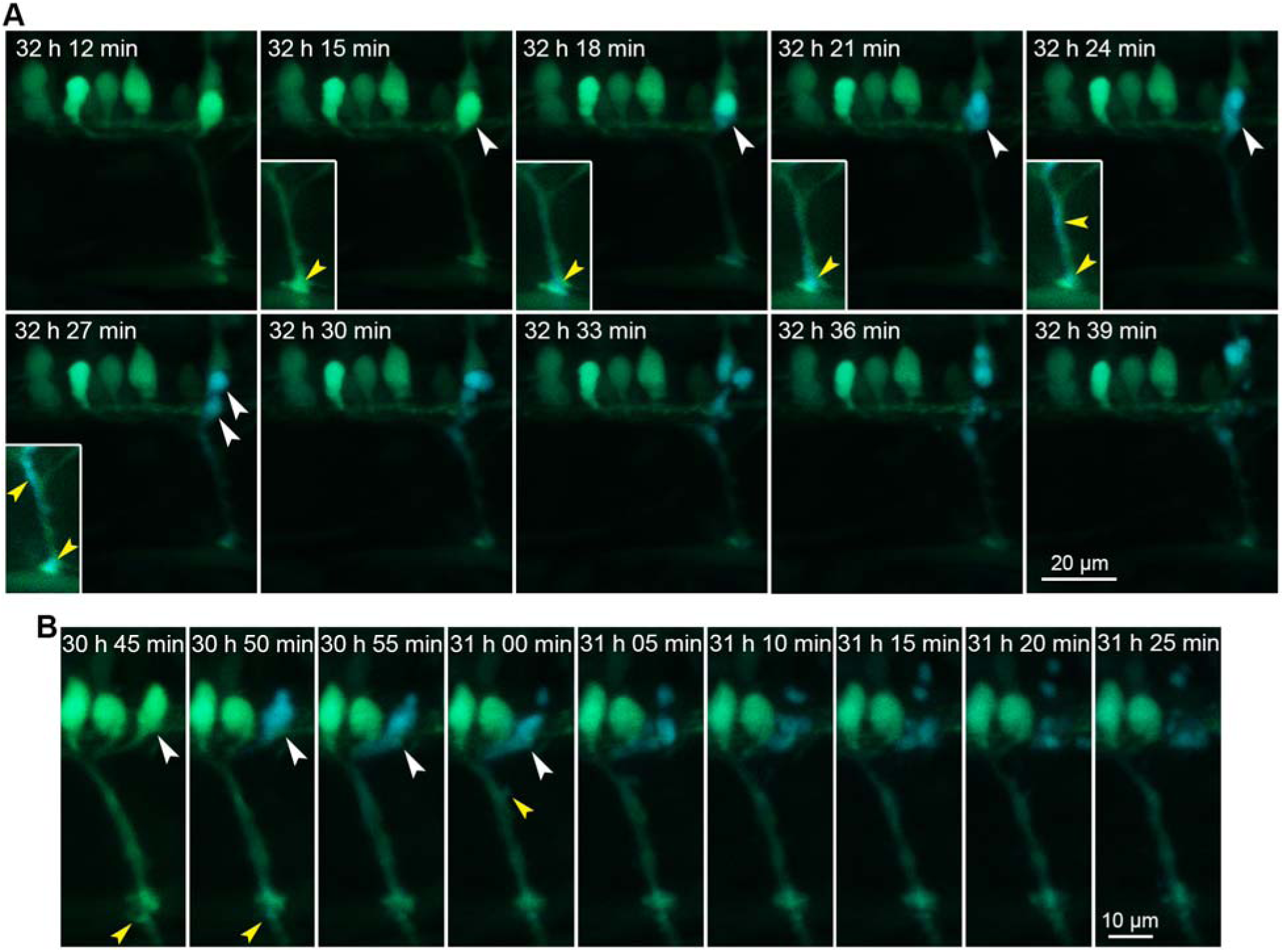
The caspase-3 activation occurred quickly and almost at the same time between the cell body and axon in apoptotic motor neurons. (**A**) Tracking of motor neuron apoptosis during zebrafish development. During tracking, one motor neuron changed from green to blue. The cell body is indicated with white arrowheads. The axon is enlarged for a better illustration and indicated with yellow arrowheads. The developmental time is indicated in each image. (**B**) Tracking of the apoptotic process in another motor neuron during zebrafish development. The cell body is indicated with white arrowheads, and the axon is indicated with yellow arrowheads. The developmental time is indicated in each image.

The time-lapse imaging of another apoptotic motor neuron showed similar results (Figure 3B; Video S2). The color of both the cell body and axon of the motor neuron changed from green to blue within 5 min (from 30 h 45 min to 30 h 50 min). This apoptotic motor neuron gradually degraded into apoptotic bodies in the next 15 min (from 30 h 50 min to 31 h 05 min). These images showed that the activation of caspase-3 can occur within 5-6 min in both the cell body and axon of an apoptotic motor neuron. Thus, we conclude that the caspase-3 activation in the cell body and axon of a dying motor neuron occurred almost at the same time.

### Only a small percentage of motor neurons died during zebrafish early development

In the last several decades, it has been widely accepted that during the embryonic development of vertebrates, massive neuronal cells in various regions of the nervous system will die. (Castillo-Ruiz, Hite, Yakout, Rosen, & Forger, 2020; Dekkers, Nikoletopoulou, & Barde, 2013; Kandel, Schwartz, Jessell, Siegelbaum, & Hudspeth, 2012; Lossi & Merighi, 2003; Rubin, 1997).

Regarding motor neuron development, chick embryos have served as the most studied models. Previous studies showed that half of motor neurons in the lateral motor column of the spinal cord died between day 5 to day 10 of incubation during the development of chick embryos (Caldero et al., 1998; Hamburger, 1975; Oppenheim, 1981; Weill, 1991). In addition to the motor neurons in the lateral motor column, a large number of motor neurons in other parts of the spinal cord also die during embryonic development of chicks (Yamamoto & Henderson, 1999). Similar motor neuron death rates were also obtained in mice (E11.5-E15.5), rats (E15-E18), and humans (11-25 weeks of gestation) during early development (Forger & Breedlove, 1987; Harris & McCaig, 1984; Raoul, Henderson, & Pettmann, 1999; Weill, 1991; Yamamoto & Henderson, 1999).

Although the motor neuron death time windows vary in different species, the common feature of these time windows is that they are all the developmental periods when motor neurons contact with muscle cells. The contact between zebrafish motor neurons and muscle cells occurs before 72 hpf (Kaslin & Ganz, 2020; Myers, Eisen, & Westerfield, 1986; Ott, Diekmann, Stuermer, & Bastmeyer, 2001; Panzer et al., 2005; Schmidt, Strähle, & Scholpp, 2013). Most organs of zebrafish form before 72 hpf, and they complete hatching before 72 hpf. Food-seeking and active avoidance behaviors also start at 72 hpf (Kimmel, Ballard, Kimmel, Ullmann, & Schilling, 1995).

To cover the potential time window of motor neuron death, we performed the noninvasive FRET imaging as long as possible: from 24 hpf to 240 hpf. After 240 hpf, the transparency of zebrafish body decreased dramatically, which made optical imaging quite difficult. Then, we counted the number of apoptotic motor neurons in the whole spinal cord from 24 hpf to 240 hpf. The data showed that the apoptosis of motor neurons mainly occurred between 24−48 hpf and peaked between 30−36 hpf (Figure 4A, B). Unexpectedly, only a tiny portion (1.6% at 30 hpf and 1.0% at 36 hpf) of the motor neurons underwent apoptosis (Figure 4C), which is much lower than the death rates in other animal models and humans.

**Figure 4.**
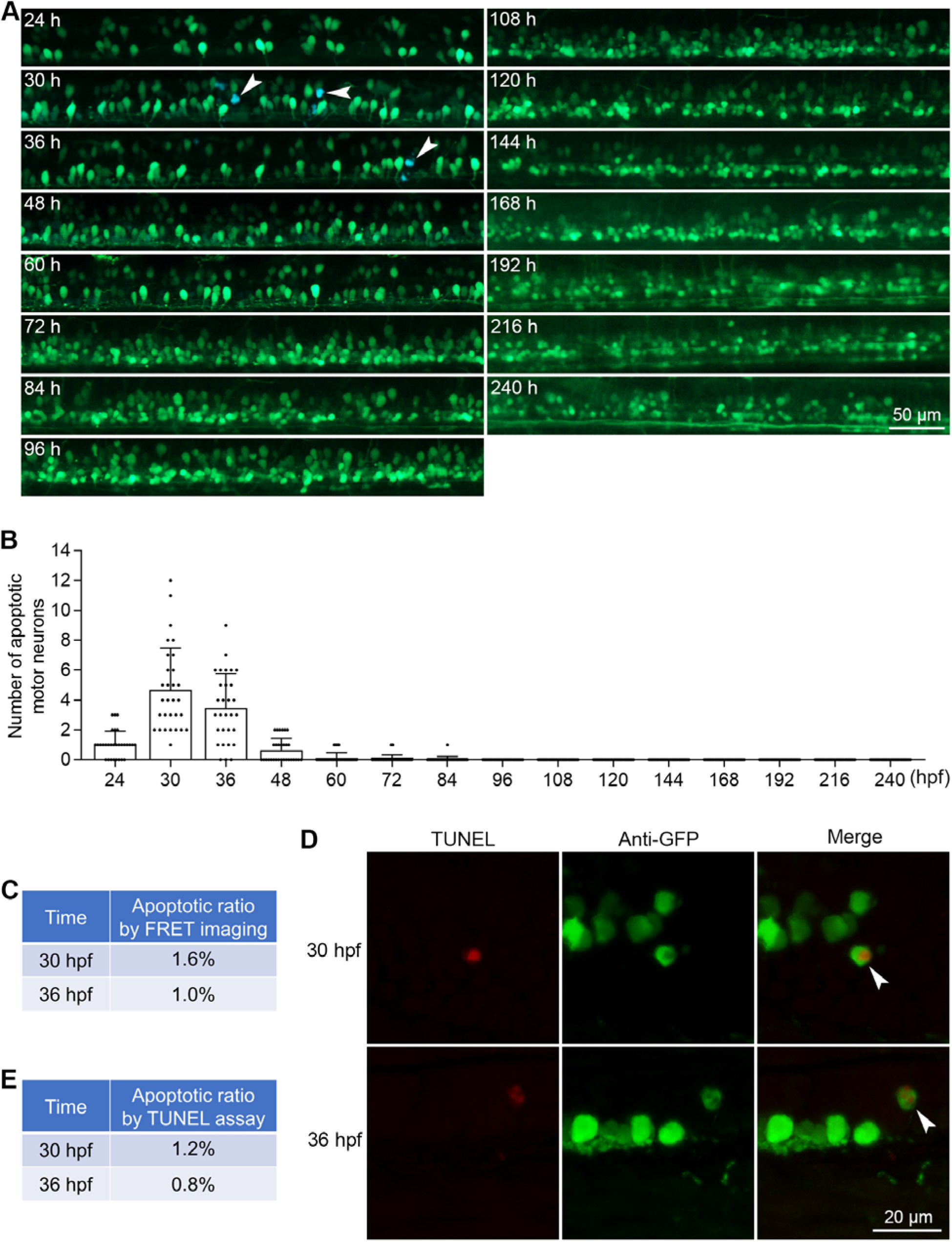
Only a small percentage of motor neurons died during zebrafish early development. (**A**) The tracking of motor neuron apoptosis by FRET imaging during zebrafish early development. White arrowheads indicate apoptotic motor neurons. (**B**) The quantified results show the number of apoptotic motor neurons at each time point during zebrafish early development (n = 30 zebrafish for each time point). (**C**) The percentages of apoptotic motor neurons among total motor neurons at 30 hpf and 36 hpf. (**D**) TUNEL assays showing apoptotic motor neurons at 30 hpf and 36 hpf. Immunofluorescence staining with an anti-GFP antibody was used to indicate motor neurons expressing sensor C3 proteins. White arrowheads indicate TUNEL-positive motor neurons. (**E**) The percentages of apoptotic motor neurons detected by TUNEL assays.

To validate these results, we performed terminal deoxynucleotidyl transferase dUTP nick end labeling (TUNEL) assays in zebrafish frozen sections. The motor neurons of the sensor zebrafish were identified by immunofluorescence staining using an anti-GFP antibody plus a secondary antibody conjugated with a green dye, and the apoptotic motor neurons were detected by red TUNEL signals (Figure 4D). We analyzed 1,045 motor neurons at 30 hpf, 13 of which were TUNEL-positive, with an apoptotic rate of 1.2%. We also analyzed 652 motor neurons at 36 hpf, 5 of which were TUNEL-positive, with an apoptotic rate of 0.8% (Figure 4E). These results showed that the low apoptotic rates detected by sensor C3-based FRET imaging could be confirmed by the conventional TUNEL assays.

### Most dead motor neurons were not colocalized with macrophages

Thus far, we have shown that sensor zebrafish can be used to detect motor neuron apoptosis. The subsequent question is how the apoptotic bodies of these dead motor neurons are removed.

Macrophages are the major phagocytes to engulf apoptotic bodies in animals (Nagata & Segawa, 2021); therefore, we focused on macrophages. We fully utilized the advantages of sensor zebrafish, which label motor neurons with green fluorescence, and another transgenic zebrafish line, Tg(*mpeg1*:mCherry), which labels macrophages with a red fluorescent protein (Ellett, Pase, Hayman, Andrianopoulos, & Lieschke, 2011). Live imaging showed that red macrophages distributed everywhere in the zebrafish body at 36 hpf (Figure 5A). Interestingly, after examining 164 apoptotic motor neurons from Tg(*mnx1*:sensor C3) zebrafish, we only found one case of macrophage colocalization with a dead motor neuron, in which the macrophage indicated with a yellow arrowhead was colocalized with the apoptotic bodies of the axon part in the somite, which were indicated with a white arrowhead (Figure 5B). The colocalization percentage was only 0.6% (Figure 5C). These results indicated that the apoptotic bodies of dead motor neurons in the spinal cord might not be cleared by macrophages.

**Figure 5.**
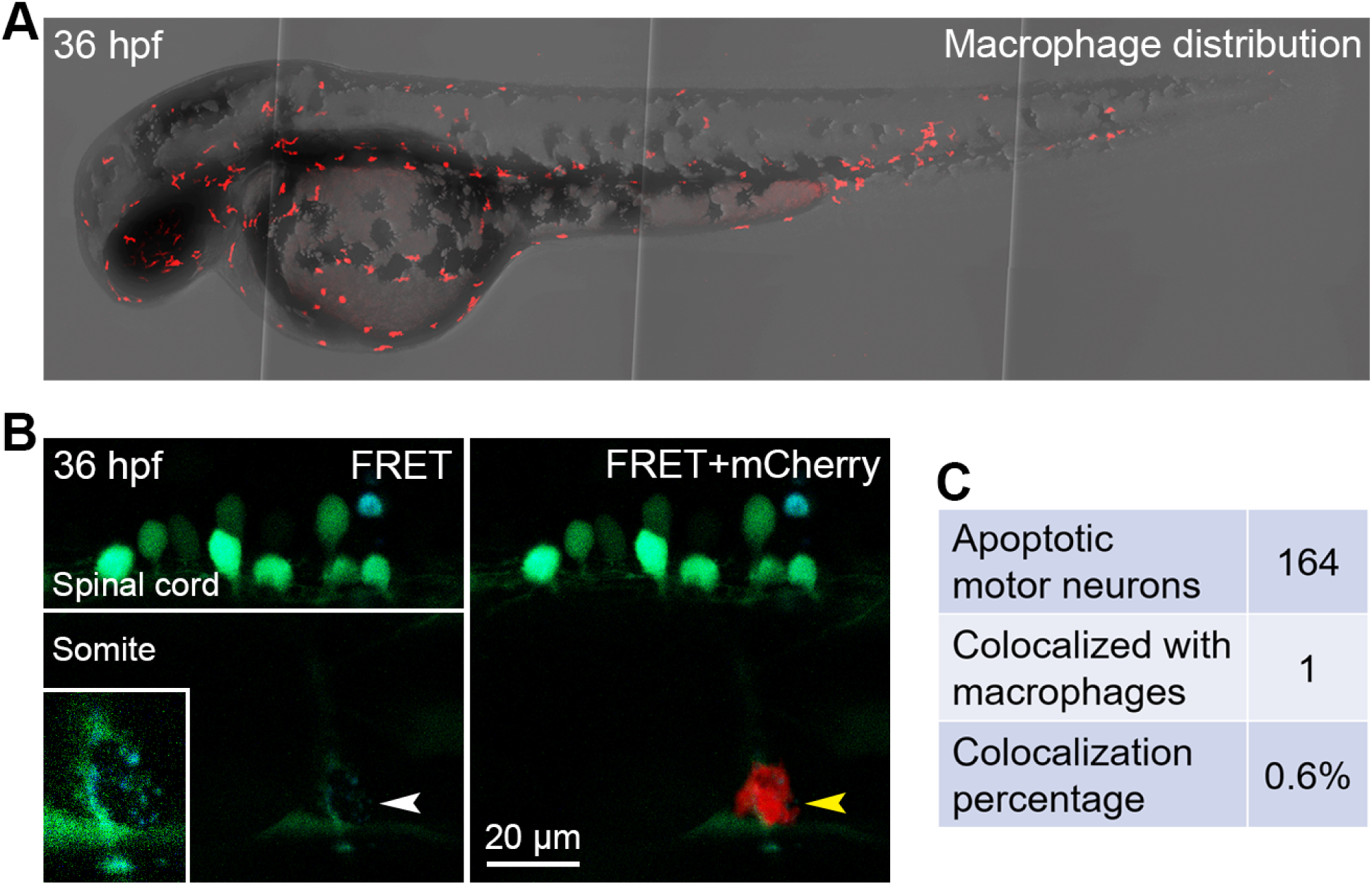
Most dead motor neurons were not colocalized with macrophages. (**A**) Live imaging showing the distribution of macrophages in zebrafish body at 36 hpf. (**B**) An example showing the colocalization between apoptotic bodies derived from the axon of a motor neuron and a macrophage in a somite at 36 hpf. The apoptotic bodies are indicated with a white arrowhead and enlarged in the box in the left image. The colocalization is indicated with a yellow arrowhead in the right image. The anatomical structures are indicated. (**C**) The percentage of apoptotic motor neurons colocalized with macrophages. The total number of apoptotic motor neurons observed and the number of apoptotic motor neurons colocalized with macrophages are listed.

## Discussion

By expressing FRET-based sensor C3 in zebrafish, for the first time, we were able to monitor the whole process of caspase-3 activation in single motor neurons and the subsequent breakdown of these dead neurons into apoptotic bodies during early development. We found that caspase-3 was activated within 5-6 min in an individual apoptotic motor neuron, while the caspase-3 in the rest of the neurons in the same bundle was not activated. More importantly, within the same motor neuron, although the cell body is located in the spinal cord and the axon is extended into the muscle tissue, the caspase-3 activation occurred almost at the same time of observation. These data indicate a synchronized activation pattern of caspase-3 in apoptotic motor neurons.

Previous studies showed that some variable primary (VaP) motor neurons died before 36 hpf during zebrafish early development (Eisen, 1992; Eisen, Pike, & Romancier, 1990). Our study examined the overall motor neuron apoptosis regardless of the causes and locations, so we did not limit our observation to VaP motor neurons. The dead motor neurons in panels A, B, and D of Figure 2 are very likely VaP motor neurons. Besides VaP motor neurons, sensor zebrafish can also capture the death of other types of motor neurons. The reasons are as follows: (1) VaP primary motor neurons die before 36 hpf, but our study found motor neurons died after 36 hpf and even at 84 hpf (Figure 4B). (2) The position of the VaP motor neuron is together with that of the caudal primary motor neuron, that is, at the caudal region of the motor neuron cluster.

Although it’s rare, we did observe the death of motor neurons in the rostral region of the motor neuron cluster (Figure 2C). (3) There is only one or zero VaP motor neuron in each motor neuron cluster. Although our data showed that usually one motor neuron died in each cluster, we did observe that sometimes more than one motor neuron died in the motor neuron cluster (Figure 2C).

It is generally accepted that during the development of vertebrates, approximately half of the motor neurons are removed by apoptosis. However, our results showed that motor neurons were not prone to death during zebrafish early development: only a small percentage of motor neurons died of apoptosis during the early development of zebrafish. This large discrepancy may be the unique properties of zebrafish compared with chicks, mice, rats, and humans, which could be further investigated. In our manuscript, we tracked the motor neuron death in live zebrafish until 240 hpf, which was the longest time window we could achieve. But there was still a possibility that zebrafish motor neurons might die after 240 hpf. Another possibility is that the limitations of conventional apoptosis detection methods, such as the TUNEL assay (Loo, 2011), nuclear morphology staining (H. Li et al., 2000), and caspase-3 staining (Wang et al., 2010), may potentially exaggerate the number of apoptotic cells in tissue sections. For example, the specificity of antibodies for apoptotic markers or the quality of other reagents in experiments may not be good enough, which can result in false positive apoptotic signals. Besides, these staining methods have difficulties in distinguishing neuronal cells and nearby glial cells; thus, dead glial cells may also be counted as dead neuronal cells.

The removal of dead cells by phagocytes can effectively reduce the occurrence of inflammation, which is very important for animals (Arandjelovic & Ravichandran, 2015). Our study showed that macrophages, although serving as professional phagocytes, were not colocalized with the apoptotic motor neurons. Thus, some other cells in the spinal cord may help to engulf the dead cells. For example, a recent zebrafish study reported that neural crest cells can help remove apoptotic cells in the nervous system during development (Zhu et al., 2019). In the future, we will investigate whether red fluorescent protein-labeled neural crest cells could engulf dead motor neurons in our sensor zebrafish.

During the past several decades, the noninvasive tracking of neuronal cell apoptosis under physiological conditions in live animals remains challenging. Since the development of fluorescent proteins, several FRET-based protein biosensors have been designed for the detection of apoptosis in live cells cultured in petri dishes (Ai, Hazelwood, Davidson, & Campbell, 2008; Albeck et al., 2008; Karasawa, Araki, Nagai, Mizuno, & Miyawaki, 2004; Mahajan, Harrison-Shostak, Michaux, & Herman, 1999; Rehm et al., 2002; Tyas, Brophy, Pope, Rivett, & Tavaré, 2000; Xu et al., 1998). However, unlike the good performances of these biosensors *in vitro*, the *in vivo* applications of these biosensors have been unsatisfactory because of their small dynamic ranges and the complex optical environments in live animals (Andrews et al., 2016; Takemoto et al., 2007; Takemoto, Nagai, Miyawaki, & Miura, 2003; Yamaguchi et al., 2011). We compared sensor C3 with other FRET-based apoptotic biosensors and found that sensor C3 can achieve a FRET effect of 4- to 5-fold, while other biosensors usually had FRET effects of less than or near 2-fold. This high FRET effect makes sensor C3 the ideal biosensor for visualizing neuronal cell apoptosis in live animals where other biosensors encounter difficulties.

In summary, our results showed that sensor zebrafish can serve as valuable tools for studying motor neuron apoptosis in live zebrafish, which could improve our understanding of neuronal cell apoptosis during normal development and in neurodegenerative diseases.

## Materials and methods Zebrafish maintenance

Zebrafish (AB strain) were raised at 28 °C in a ZebTEC system. The photoperiod was 14 h of light and 10 h of darkness. Zebrafish were fed with artemia twice a day. The experiments in this study were approved by the Animal Research Ethics Committee of the University of Macau (Protocol ID: UMARE-032-2016).

## Generation of sensor zebrafish

The zebrafish genome was extracted using the DNeasy Blood & Tissue Kit (Qiagen, 69504). A zebrafish motor neuron-specific promoter *mnx1* was amplified by polymerase chain reaction (forward primer, ACGCGTCGACGAATTCATTTAAATTAGCCTGGCATC, reverse primer, CCCACCGGTCTGGCCCACCTCACAAACAGATTA). The promoter sequence and sensor C3 gene were cloned into the same plasmid backbone to generate the transgenic vector, named pSK- *mnx1*-C3. *Tol2* mRNA was obtained by *in vitro* transcription using the mMESSAGE mMACHINE SP6 transcription kit (Ambion, AM1340). The transgenic vector containing the sensor C3 gene (final concentration, 50 ng/μL) and *Tol2* mRNA (final concentration, 100 ng/μL) were mixed and injected (2-4 nL) into newly fertilized zebrafish eggs using an MPPI-3 pressure injector. The zebrafish embryos with green fluorescence indicating sensor C3 expression were selected under a stereomicroscope with a GFP filter. These positive zebrafish were cultured to sexual maturity and crossed with wild-type zebrafish to obtain transgenic sensor zebrafish.

## *In vivo* FRET imaging of neuronal cell apoptosis

Sensor zebrafish embryos or larvae were anesthetized with 0.016% tricaine (Sigma-Aldrich, E10521) solution. Anesthetized zebrafish were embedded in low melting point agarose gel (Promega, V2111) in a confocal dish. The zebrafish were then imaged using a Carl Zeiss LSM 710 or LSM880 confocal laser scanning microscope. For FRET imaging, a 458 nm laser was used to excite sensor C3 proteins, and the emissions were collected simultaneously in two channels: 460-500 nm was collected as the CFP channel, designated as blue, and 520-550 nm was collected as the YFP channel, designated as green. Then, the CFP and YFP images were merged to generate the FRET images. In live neuronal cells, sensor C3 emitted green fluorescence when CFP was excited because of the energy transfer from CFP to YFP, while in apoptotic neuronal cells, activated caspase-3 proteins cleaved sensor C3 at the DEVD site, and the FRET effect was abolished. Thus, sensor C3 emitted blue fluorescence when CFP was excited under the same condition.

## Immunofluorescence and TUNEL assay

Tg(*mnx1*:sensor C3) zebrafish were fixed in 4% paraformaldehyde at 4 °C for 12 h. The fixed zebrafish were dehydrated in 30% sucrose at 4 °C for 48 h. The zebrafish were then embedded in Shandon Cryomatrix embedding resin (Thermo Scientific, 6769006), and frozen sections were cut at 25 µm using a Leica CM3050S cryostat. The sections were washed with phosphate-buffered saline (PBS) three times to remove the embedding resin and then were blocked with 1% bovine serum albumin with 0.1% Triton X-100 in PBS for 1 h at room temperature (RT). Then, the sections were coated with anti-GFP primary antibody (Cell Signaling, 2956, 1:400) for 2 h at RT. After that, the sections were washed with PBS three times and then coated with Alexa Fluor 488-conjugated goat anti-rabbit secondary antibody (Thermo Scientific, A-11034, 1:200) for 1 h at RT. The sections were washed with PBS three times. The terminal deoxynucleotidyl transferase and Cy3-dUTP labeling solution in the TUNEL kit (Beyotime, C1089) were mixed.

The sections were then incubated with this reaction mixture for 1 h at 37 °C. After the incubation, the sections were washed with PBS three times and then mounted with Mowiol mounting medium (Millipore, 475904). The sections were imaged with a Carl Zeiss LSM880 confocal laser scanning microscope with 488 nm and 561 nm lasers.

## *In vivo* imaging of macrophages

Tg(*mnx1*:sensor C3) zebrafish were crossed with Tg(*mpeg1*:mCherry) zebrafish to get offspring that neuronal cells and macrophages were labeled with green and red, respectively. The zebrafish were then imaged using a Carl Zeiss LSM880 confocal laser scanning microscope. FRET imaging was applied to detect apoptotic neuronal cells. At the same time, a 561 nm laser was used to excite the mCherry proteins for the visualization of macrophages.

## Statistical analysis

All of the data are expressed as the means ± SD. Statistical significance was judged by one- or two-way ANOVA using GraphPad Prism 7 software.

## Supplementary materials

Supplementary materials can be found online.

## Acknowledgments

The authors acknowledge the great support of the Animal Research Core and Biological Imaging and Stem Cell Core of the Faculty of Health Sciences at the University of Macau. The transgenic zebrafish line Tg(*mpeg1*:mCherry) was a gift from Dr. Kun Wu. This work was financially supported by the Multi-Year Research Grant of University of Macau (File no. MYRG2020- 00121-FHS and MYRG2022-00025-FHS), the Science and Technology Development Fund (FDCT) of Macao (File no. 0147/2020/A3, 044/2021/APD and 0004/2021/AKP), Ministry of Education Frontiers Science Centre for Precision Oncology (File no. SP2021-00001-FSCPO and SP2023-00001-FSCPO) and an FHS Internal Project Grant of the University of Macau.

## Author contributions

H.J. and K.Q.L. designed the experiments. H.J. and H.Y. performed the experiments. H.J. and K.Q.L. analyzed the data. H.J. prepared the manuscript. K.Q.L. revised the manuscript. K.Q.L. supervised the study.

## Data and materials availability

All data needed to evaluate the conclusions in the paper are present in the paper and/or the Supplementary materials.

## References

1. Ai, H.-w., Hazelwood, K. L., Davidson, M. W., & Campbell, R. E. (2008). Fluorescent protein FRET pairs for ratiometric imaging of dual biosensors. Nat. Methods, 5(5), 401–403.

2. Albeck, J. G., Burke, J. M., Aldridge, B. B., Zhang, M., Lauffenburger, D. A., & Sorger, P. K. (2008). Quantitative analysis of pathways controlling extrinsic apoptosis in single cells. Mol. Cell, 30(1), 11–25.

3. Anand, P., Fu, A., Teoh, S. H., & Luo, K. Q. (2015). Application of a fluorescence resonance energy transfer (FRET)[based biosensor for detection of drug[induced apoptosis in a 3D breast tumor model. Biotechnol. Bioeng., 112(8), 1673–1682.

4. Andrews, N., Ramel, M. C., Kumar, S., Alexandrov, Y., Kelly, D. J., Warren, S. C., et al. (2016). Visualising apoptosis in live zebrafish using fluorescence lifetime imaging with optical projection tomography to map FRET biosensor activity in space and time. J. Biophotonics, 9(4), 414–424.

5. Arandjelovic, S., & Ravichandran, K. S. (2015). Phagocytosis of apoptotic cells in homeostasis. Nat.Immunol., 16(9), 907–917.

6. Arkhipova, V., Wendik, B., Devos, N., Ek, O., Peers, B., & Meyer, D. (2012). Characterization and regulation of the hb9/mnx1 beta-cell progenitor specific enhancer in zebrafish. Dev. Biol., 365(1), 290–302.

7. Blanquie, O., Yang, J.-W., Kilb, W., Sharopov, S., Sinning, A., & Luhmann, H. J. (2017). Electrical activity controls area-specific expression of neuronal apoptosis in the mouse developing cerebral cortex. eLife, 6, e27696.

8. Butterfield, D. A., & Halliwell, B. (2019). Oxidative stress, dysfunctional glucose metabolism and Alzheimer disease. Nat. Rev. Neurosci., 20(3), 148–160.

9. Caldero, J., Prevette, D., Mei, X., Oakley, R. A., Li, L., Milligan, C., et al. (1998). Peripheral target regulation of the development and survival of spinal sensory and motor neurons in the chick embryo. J. Neurosci., 18(1), 356–370.

10. Castillo-Ruiz, A., Hite, T. A., Yakout, D. W., Rosen, T. J., & Forger, N. G. (2020). Does birth trigger cell death in the developing brain? eNeuro, 7(1), 1–11.

11. Dekkers, M. P., Nikoletopoulou, V., & Barde, Y.-A. (2013). Death of developing neurons: New insights and implications for connectivity. J. Cell Biol., 203(3), 385–393.

12. Eisen, J. S. (1992). The role of interactions in determining cell fate of two identified motoneurons in the embryonic zebrafish. Neuron, 8(2), 231–240.

13. Eisen, J. S., Pike, S., & Romancier, B. (1990). An identified motoneuron with variable fates in embryonic zebrafish. J. Neurosci., 10(1), 34–43.

14. Ellett, F., Pase, L., Hayman, J. W., Andrianopoulos, A., & Lieschke, G. J. (2011). mpeg1 promoter transgenes direct macrophage-lineage expression in zebrafish. Blood, 117(4), e49–e56.

15. Flanagan-Steet, H., Fox, M. A., Meyer, D., & Sanes, J. R. (2005). Neuromuscular synapses can form in vivo by incorporation of initially aneural postsynaptic specializations. Development.

16. Forger, N. G., & Breedlove, S. M. (1987). Motoneuronal death during human fetal development. J. Comp. Neurol., 264(1), 118–122.

17. Fu, A., Peh, Y. M., Ngan, W., Wei, N., & Luo, K. Q. (2018). Rapid identification of antimicrometastases drugs using integrated model systems with two dimensional monolayer, three dimensional spheroids, and zebrafish xenotransplantation tumors. Biotechnol. Bioeng., 115(11), 2828–2843.

18. Fuchs, Y., & Steller, H. (2011). Programmed cell death in animal development and disease. Cell, 147(4), 742–758.

19. Hamburger, V. (1975). Cell death in the development of the lateral motor column of the chick embryo. Journal of Comparative Neurology, 160(4), 535–546.

20. Hao, M., Huang, B., Wu, R., Peng, Z., & Luo, K. Q. (2023). The Interaction between Macrophages and Triple[negative Breast Cancer Cells Induces ROS[Mediated Interleukin 1α Expression to Enhance Tumorigenesis and Metastasis. Advanced Science, 2302857.

21. Harris, A., & McCaig, C. (1984). Motoneuron death and motor unit size during embryonic development of the rat. Journal of Neuroscience, 4(1), 13–24.

22. Heck, N., Golbs, A., Riedemann, T., Sun, J.-J., Lessmann, V., & Luhmann, H. J. (2008). Activity-dependent regulation of neuronal apoptosis in neonatal mouse cerebral cortex. Cereb. Cortex, 18(6), 1335–1349.

23. Hernandez-Baltazar, D., Mendoza-Garrido, M. E., & Martinez-Fong, D. (2013). Activation of GSK-3β and caspase-3 occurs in Nigral dopamine neurons during the development of apoptosis activated by a striatal injection of 6-hydroxydopamine. PLoS One, 8(8), e70951.

24. Hollville, E., Romero, S. E., & Deshmukh, M. (2019). Apoptotic cell death regulation in neurons. FEBS J., 286(17), 3276–3298.

25. Jao, L.-E., Appel, B., & Wente, S. R. (2012). A zebrafish model of lethal congenital contracture syndrome 1 reveals Gle1 function in spinal neural precursor survival and motor axon arborization. Development, 139(7), 1316–1326.

26. Jia, H., & Luo, K. Q. (2021). Fluorescence resonance energy transfer-based sensor zebrafish for detecting toxic agents with single-cell sensitivity. J. Hazard. Mater., 408, 124826.

27. Jia, H., Song, Y., Huang, B., Ge, W., & Luo, K. Q. (2020). Engineered sensor zebrafish for fast detection and real-time tracking of apoptosis at single-cell resolution in live animals. ACS Sens., 5(3), 823–830.

28. Kandel, E. R., Schwartz, J. H., Jessell, T. M., Siegelbaum, S. A., & Hudspeth, A. J. (2012). Principles of Neural Science, Fifth Edition: McGraw-Hill New York.

29. Karasawa, S., Araki, T., Nagai, T., Mizuno, H., & Miyawaki, A. (2004). Cyan-emitting and orange-emitting fluorescent proteins as a donor/acceptor pair for fluorescence resonance energy transfer. Biochem. J., 381(1), 307–312.

30. Kaslin, J., & Ganz, J. (2020). Zebrafish nervous systems. In The zebrafish in biomedical research (pp.181–189): Elsevier.

31. Kimmel, C. B., Ballard, W. W., Kimmel, S. R., Ullmann, B., & Schilling, T. F. (1995). Stages of embryonic development of the zebrafish. Dev. Dynam., 203(3), 253–310.

32. Li, H., Kolluri, S. K., Gu, J., Dawson, M. I., Cao, X., Hobbs, P. D., et al. (2000). Cytochrome c release and apoptosis induced by mitochondrial targeting of nuclear orphan receptor TR3. Science, 289(5482), 1159–1164.

33. Li, K., Wu, R., Zhou, M., Tong, H., & Luo, K. Q. (2021). Desmosomal proteins of DSC2 and PKP1 promote cancer cells survival and metastasis by increasing cluster formation in circulatory system. Sci. Adv., 7(40), eabg7265.

34. Loo, D. T. (2011). In situ detection of apoptosis by the TUNEL assay: an overview of techniques. In DNA damage detection in situ, ex vivo, in vivo (Vol. 682, pp. 3–13): Humana Press.

35. Lossi, L., & Merighi, A. (2003). In vivo cellular and molecular mechanisms of neuronal apoptosis in the mammalian CNS. Prog. Neurobiol., 69(5), 287–312.

36. Luo, K. Q., Vivian, C. Y., Pu, Y., & Chang, D. C. (2001). Application of the fluorescence resonance energy transfer method for studying the dynamics of caspase-3 activation during UV-induced apoptosis in living HeLa cells. Biochem. Biophys. Res. Commun., 283(5), 1054–1060.

37. Mahajan, N. P., Harrison-Shostak, D. C., Michaux, J., & Herman, B. (1999). Novel mutant green fluorescent protein protease substrates reveal the activation of specific caspases during apoptosis. Chem. Biol., 6(6), 401–409.

38. Michel, P. P., Hirsch, E. C., & Hunot, S. (2016). Understanding dopaminergic cell death pathways in Parkinson disease. Neuron, 90(4), 675–691.

39. Moujalled, D., Strasser, A., & Liddell, J. R. (2021). Molecular mechanisms of cell death in neurological diseases. Cell Death Differ., 28(7), 2029–2044.

40. Myers, P. Z., Eisen, J. S., & Westerfield, M. (1986). Development and axonal outgrowth of identified motoneurons in the zebrafish. J. Neurosci., 6(8), 2278–2289.

41. Nagata, S., & Segawa, K. (2021). Sensing and clearance of apoptotic cells. Curr. Opin. Immunol., 68, 1–8.

42. Oppenheim, R. W. (1981). Cell death of motoneurons in the chick embryo spinal cord. V. Evidence on the role of cell death and neuromuscular function in the formation of specific peripheral connections. Journal of Neuroscience, 1(2), 141–151.

43. Ott, H., Diekmann, H., Stuermer, C. A., & Bastmeyer, M. (2001). Function of Neurolin (DM-GRASP/SC-1) in guidance of motor axons during zebrafish development. Dev. Biol., 235(1), 86–97.

44. Panzer, J. A., Gibbs, S. M., Dosch, R., Wagner, D., Mullins, M. C., Granato, M., et al. (2005). Neuromuscular synaptogenesis in wild-type and mutant zebrafish. Dev. Biol., 285(2), 340–357.

45. Raoul, C., Henderson, C. E., & Pettmann, B. (1999). Programmed cell death of embryonic motoneurons triggered through the Fas death receptor. J. Cell Biol., 147(5), 1049–1062.

46. Rehm, M., Dussmann, H., Janicke, R. U., Tavare, J. M., Kogel, D., & Prehn, J. H. (2002). Single-cell fluorescence resonance energy transfer analysis demonstrates that caspase activation during apoptosis is a rapid process: role of caspase-3. J. Biol. Chem., 277(27), 24506–24514.

47. Rosenberg, A. F., Wolman, M. A., Franzini-Armstrong, C., & Granato, M. (2012). In vivo nerve– macrophage interactions following peripheral nerve injury. J. Neurosci., 32(11), 3898–3909.

48. Rubin, L. L. (1997). Neuronal cell death: when, why and how. Br. Med. Bull., 53(3), 617–631.

49. Schmidt, R., Strähle, U., & Scholpp, S. (2013). Neurogenesis in zebrafish–from embryo to adult. J Neural Development, 8(1), 1–13.

50. Stil, A., & Drapeau, P. (2016). Neuronal labeling patterns in the spinal cord of adult transgenic Zebrafish. Dev. Neurobiol., 76(6), 642–660.

51. Takemoto, K., Kuranaga, E., Tonoki, A., Nagai, T., Miyawaki, A., & Miura, M. (2007). Local initiation of caspase activation in Drosophila salivary gland programmed cell death in vivo. Proc. Natl. Acad. Sci. U. S. A., 104(33), 13367–13372.

52. Takemoto, K., Nagai, T., Miyawaki, A., & Miura, M. (2003). Spatio-temporal activation of caspase revealed by indicator that is insensitive to environmental effects. J. Cell Biol., 160(2), 235–243.

53. Tian, H., Ip, L., Luo, H., Chang, D. C., & Luo, K. Q. (2007). A high throughput drug screen based on fluorescence resonance energy transfer (FRET) for anticancer activity of compounds from herbal medicine. Br. J. Pharmacol., 150(3), 321–334.

54. Tyas, L., Brophy, V. A., Pope, A., Rivett, A. J., & Tavaré, J. M. (2000). Rapid caspase -3 activation during apoptosis revealed using fluorescence-resonance energy transfer. EMBO Rep., 1(3), 266–270.

55. Wang, Q., Frolova, A. I., Purcell, S., Adastra, K., Schoeller, E., Chi, M. M., et al. (2010). Mitochondrial dysfunction and apoptosis in cumulus cells of type I diabetic mice. PloS One, 5(12), e15901.

56. Weill, C. (1991). Somatostatin (SRIF) prevents natural motoneuron cell death in embryonic chick spinal cord. Dev. Neurosci., 13(6), 377–381.

57. Wu, R., Li, K., Yuan, M., & Luo, K. Q. (2021). Nerve growth factor receptor increases the tumor growth and metastatic potential of triple-negative breast cancer cells. Oncogene, 40(12), 2165–2181.

58. Xu, X., Gerard, A. L., Huang, B. C., Anderson, D. C., Payan, D. G., & Luo, Y. (1998). Detection of programmed cell death using fluorescence energy transfer. Nucleic Acids Res., 26(8), 2034–2035.

59. Yamaguchi, Y., Shinotsuka, N., Nonomura, K., Takemoto, K., Kuida, K., Yosida, H., et al. (2011). Live imaging of apoptosis in a novel transgenic mouse highlights its role in neural tube closure. J. Cell Biol., 195(6), 1047–1060.

60. Yamamoto, Y., & Henderson, C. E. (1999). Patterns of programmed cell death in populations of developing spinal motoneurons in chicken, mouse, and rat. Dev. Biol., 214(1), 60–71.

61. Yang, H., Jia, H., Zhao, Q., & Luo, K. Q. (2022). Visualization of natural killer cell-mediated killing of cancer cells at single-cell resolution in live zebrafish. Biosens. Bioelectron., 216, 114616.

62. Zhu, Y., Crowley, S. C., Latimer, A. J., Lewis, G. M., Nash, R., & Kucenas, S. (2019). Migratory neural crest cells phagocytose dead cells in the developing nervous system. Cell, 179(1), 74–89.

